# Unraveling the Mechanistic Links Between Blood Pressure Regulation and Calcium-Magnesium Homeostasis: Insights into Hypertension, Hyperparathyroidism, and Mineral Disorders

**DOI:** 10.1101/2025.10.26.684662

**Authors:** Pritha Dutta, Anita T. Layton

## Abstract

The systems regulating blood pressure and calcium-magnesium (Ca^2+^-Mg^2+^) homeostasis are increasingly recognized to have significant, clinically relevant interactions, where alterations in one can lead to significant changes in the other. In this study, we developed a computational model integrating blood pressure regulation and Ca^2^□-Mg^2^□ homeostasis in a male rat. We simulated various conditions, including hypertension, Ca^2^□, Mg^2^□, and vitamin D_3_ deficiencies, and primary hyperparathyroidism. Simulations of hypertension, induced by various stimuli like increased renin or aldosterone secretion, demonstrated significant effects on parathyroid hormone (PTH), calcitriol, renal Ca^2^□/Mg^2^□ handling, and bone resorption. Dietary Ca^2^□, Mg^2^□, and vitamin D_3_ deficiencies elevated mean arterial pressure, with Mg^2^□ deficiency having a stronger effect. Furthermore, the model predicted that primary hyperparathyroidism elevates PTH, Ca^2^□, and calcitriol, leading to increased mean arterial pressure and bone loss. Overall, this model provides valuable insights into the mechanistic links between blood pressure regulation and Ca^2^□-Mg^2^□ homeostasis, offering insights into clinical conditions like hypertension and hyperparathyroidism.

## 1. Introduction

Hypertension is the leading cause of cardiovascular disease worldwide (1), and its prevalence has been rising in recent years due to factors such as an aging population and lifestyle changes, including decreased physical activity and the widespread consumption of Western diets (2). Furthermore, a significant portion of the global population fails to meet the recommended dietary intake of magnesium (Mg^2^□) and calcium (Ca^2^□) (3,4). For example, the standard diet in the United States provides only about 50% of the recommended daily intake of Mg^2^□ (5), and the 2017-2018 National Health and Nutrition Examination Survey (NHANES) found that nearly half of the US population did not meet their estimated average requirements for Ca^2+^ (6). Additionally, approximately 100,000 people in the US are diagnosed with primary hyperparathyroidism each year (7). Given these factors, it is essential to understand how blood pressure regulation and Ca^2+^-Mg^2+^ homeostasis are affected by these conditions.

The systems regulating blood pressure and Ca^2+^-Mg^2+^ homeostasis are increasingly recognized to have significant, clinically relevant interactions (8–10). A key component of blood pressure regulation is the renin-angiotensin-aldosterone system (RAAS). Ca^2+^, Mg^2+^, and their regulatory hormones, parathyroid hormone (PTH) and calcitriol (1,25(OH)_2_D_3_; active form of vitamin D_3_), play a crucial role in regulating various elements of the RAAS (8–10). For instance, Ca^2+^ and calcitriol influence renin secretion from the juxtaglomerular (JG) cells in the kidneys (11–14). Additionally, Ca^2+^, Mg^2+^, and PTH regulate aldosterone secretion from the adrenal gland (15–19) as well as vascular resistance (20–22). Both aldosterone and angiotensin II also influence PTH secretion from the parathyroid gland (23). Furthermore, PTH and Ca^2+^ control sodium (Na^+^) reabsorption in the proximal tubule and thick ascending limb of the kidneys (24,25), which in turn affects the reabsorption of Ca^2+^ and Mg^2+^ in these nephron segments. Angiotensin II and aldosterone also indirectly influence Ca^2+^ and Mg^2+^ reabsorption in the kidneys by regulating Na^+^ reabsorption. Finally, Ca^2+^ and Mg^2+^ have opposing effects on renal sympathetic nervous activity (RSNA) (26,27). Given these multiple interconnections, a dysregulation in one system can significantly impact the other. Therefore, understanding the interaction between these systems is increasingly important for accurately assessing their impact on health.

Mechanistic modeling provides a comprehensive framework for understanding and analyzing complex physiological systems. In this study, we integrated our previously developed Ca^2+^-Mg^2+^ homeostasis model in a male rat (28) with a blood pressure regulation model in a male rat (29), quantifying the interactions between these two systems. This integrated model was then used to simulate various conditions, including hypertension, Ca^2+^, Mg^2+^, and vitamin D_3_ deficiencies, and primary hyperparathyroidism.

## 2. Methods

We have developed a model that captures the interactions between blood pressure regulation and Ca^2+^-Mg^2+^ homeostasis in a male rat. This model was developed by integrating our previously developed model of Ca^2+^ and Mg^2+^ homeostasis (28) with a blood pressure regulation model in a male rat (29). The blood pressure regulation model describes, using a large system of coupled nonlinear algebraic differential equations, the interactions among the cardiovascular system, the renal system, the renal sympathetic nervous system, and the RAAS. A schematic diagram is shown in Fig. 1 (A). Model equations and parameter values can be found in Ref. (29). The Ca^2+^ and Mg^2+^ homeostasis model consists of five compartments: plasma, parathyroid gland, intestine, kidney, and bone. The model describes, using a large system of coupled nonlinear ordinary differential equations, the exchanges of Ca^2+^, Mg^2+^, PTH, and calcitriol. A schematic diagram is shown in Fig. 1 (B). Model equations and parameter values can be found in Ref. (28).

**Figure 1:**
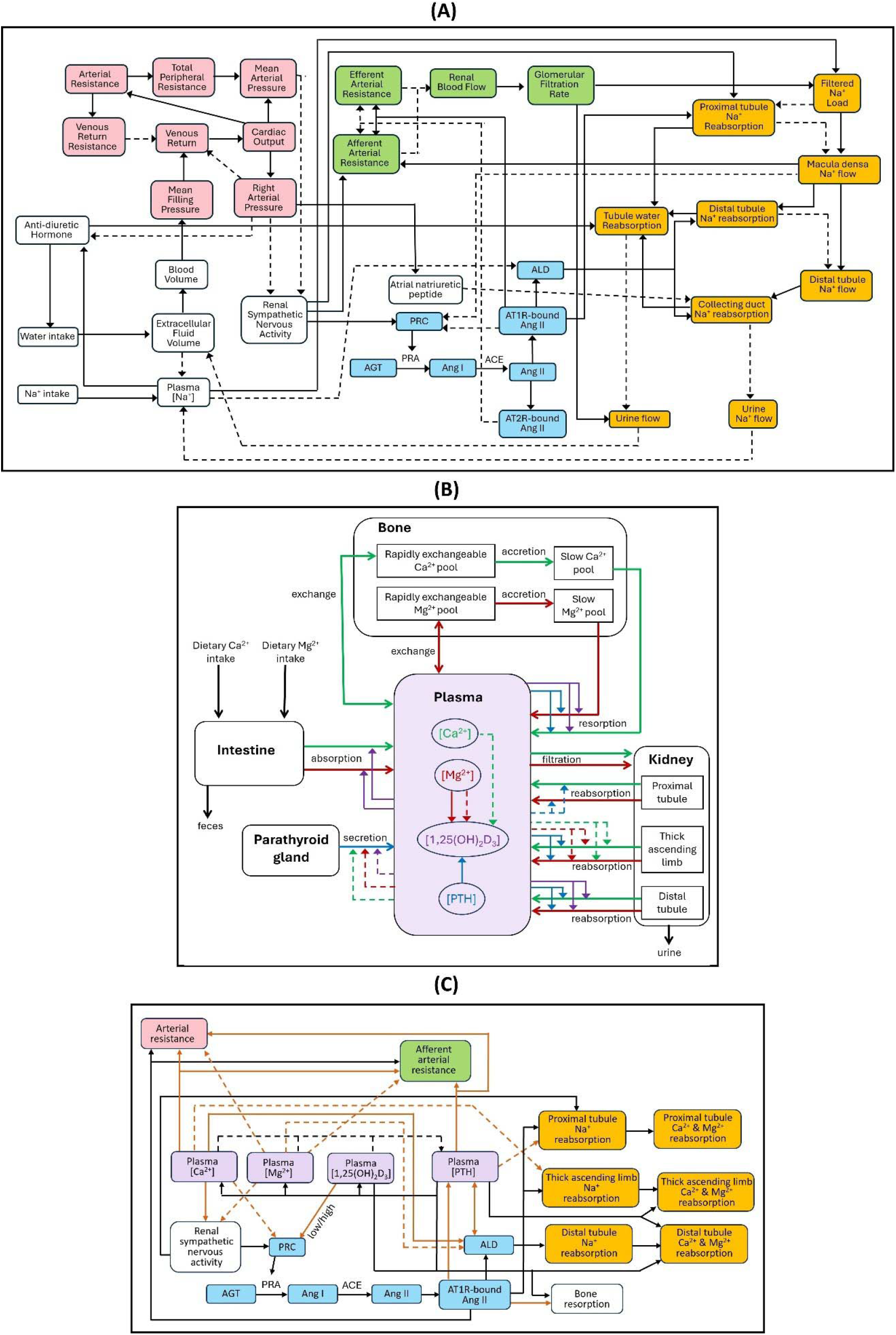
Schematic diagram of the integrated model. (A) Schematic model of blood pressure regulation. Pink nodes denote variables that describe cardiovascular function; green nodes, renal hemodynamics; yellow nodes, renal Na^+^ and fluid handling; blue nodes, the renin-angiotensin-aldosterone system. Solid arrows indicate activation and dotted arrows indicate inhibition. (B) Schematic model of Ca^2+^-Mg^2+^ homeostasis. The model consists of five compartments: plasma, intestine, kidney, parathyroid gland, and bone. Solid arrows with open arrowheads indicate fluxes, solid arrows with closed arrowheads indicate activation, and dotted arrows indicate inhibition. All arrows are color coded. Green arrows, Ca^2+^; red arrows, Mg^2+^, blue arrows, parathyroid hormone (PTH); purple arrows, 1,25(OH)_2_D_3_. (C) Schematic representation of the interactions between the models shown in (A) and (B). Mauve nodes denote plasma [Ca^2+^], [Mg^2+^], [1,25(OH)2D3], and [PTH]; yellow nodes, renal Na^+^, Ca^2+^, and Mg^2+^ handling; blue nodes, the renin-angiotensin-aldosterone system. Solid arrows indicate activation and dotted arrows indicate inhibition. The brown arrows indicate the direct links between components of the blood pressure regulation model and Ca^2+^-Mg^2+^ homeostasis model. ACE, angiotensin-converting enzyme; AGT, angiotensinogen; ALD, aldosterone; Ang I, angiotensin I; Ang II, angiotensin II; AT1R-bound Ang II, angiotensin II type 1 receptor-bound angiotensin II; AT2R-bound Ang II, angiotensin II type 2 receptor-bound angiotensin II; PRA, plasma renin activity; PRC, plasma renin concentration.

In the following sections we define the equations for the regulation of different components of the blood pressure regulation model by Ca^2+^, Mg^2+^, PTH, and 1,25(OH)_2_D_3_ and vice-versa. Figure 1 (C) provides a schematic representation of the interactions between the two systems. All new parameter values and descriptions are listed in Table 1.

**Table 1:** Description and values of new model parameters and variables at baseline for male rat.

| Symbol | Description | Value | Reference | Comments |
| --- | --- | --- | --- | --- |
| $K_{R-Ca}$ | Sensitivity of renin secretion to plasma $Ca^{2+}$ concentration | 5.4 mM | (11) | study conducted on rats |
| $v_{Ca}$ | Effect of $Ca^{2+}$ on renin secretion | 1 | | |
| $K_{R-1,25(OH)_2D_3-low}$ | Sensitivity of renin secretion to low plasma $1,25(OH)_2D_3$ concentration | 30 pM | (13) | study conducted on mice |
| $K_{R-1,25(OH)_2D_3-high}$ | Sensitivity of renin secretion to high plasma $1,25(OH)_2D_3$ concentration | 200 pM | (14) | study conducted on rats |
| $v_{1,25(OH)_2D_3}$ | Effect of $1,25(OH)_2D_3$ on renin secretion | 1 | | |
| $K_{ALD-Ca}$ | Sensitivity of aldosterone secretion to plasma Ca concentration | 2.67 mM | (15) | study conducted on rats |
| $\xi_{Ca}$ | Effect of $Ca^{2+}$ on aldosterone secretion | 1 | | |
| $K_{ALD-Mg}$ | Sensitivity of aldosterone secretion to plasma Mg concentration | 1.03 mM | (16) | study conducted on rats |
| $\xi_{Mg}$ | Effect of $Mg^{2+}$ on aldosterone secretion | 1 | | |
| $K_{ALD-PTH}$ | Sensitivity of aldosterone secretion to plasma PTH concentration | 124 pM | (37) | study conducted on rats |
| $\xi_{PTH}$ | Effect of PTH on aldosterone secretion | 1 | | |
| $K_{PTH-ALD}$ | Sensitivity of PTH secretion to aldosterone concentration | 800 ng/l | (23) | study conducted on human parathyroid cells |
| $K_{PTH-AT1R}$ | Sensitivity of PTH secretion to AT1R- bound Ang II concentration | 42 fmol/ml | (23) | |
| $\alpha_{Ca}$ | Effect of $Ca^{2+}$ on RSNA | 1 | | |
| $\alpha_{Mg}$ | Effect of $Mg^{2+}$ on RSNA | 1 | | |
| $K_{res}^{AT1R}$ | Sensitivity of bone resorption to AT1R-bound Ang II | 20 fmol/ml | | Assumed to be equal to the AT1R-bound Ang II concentration at equilibrium. |
| $K_{sod-PTH}$ | Sensitivity of $Na^+$ reabsorption in proximal tubule to PTH | 2.39 pM | | |
| $\gamma_{PTH}$ | Effect of PTH on fractional $Na^+$ reabsorption in the proximal tubule | 1 | | |
| $K_{sod-Ca}$ | Sensitivity of $Na^+$ reabsorption in thick ascending limb to $Ca^{2+}$ | 1.25 mM | (38) | |
| $\gamma_{Ca}$ | Effect of $Ca^{2+}$ on fractional $Na^+$ reabsorption in the thick ascending limb | 1 | | |
| $\Psi_{AT1R}$ | Effect of AT1R-bound Ang II on arterial resistance | 1 | | |
| $\Psi_{Ca}$ | Effect of $Ca^{2+}$ on vascular resistance | 1 | | |
| $\Psi_{Mg}$ | Effect of $Mg^{2+}$ on vascular resistance | 1 | | |
| $\Psi_{PTH}$ | Effect of PTH on vascular resistance | 1 | | |

### 2.1 Renin secretion

Renin is secreted by the juxtaglomerular (JG) cells in the kidneys. Renin secretion is given by

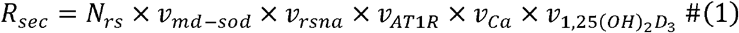

where *N*_*rs*_ represents the normalized renin secretion, and *v*_*md*−*sod*_, *v*_*rsna*_,*v*_*AT*1*R*_, *v*_*Ca*,_ and 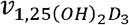 represent the effect of tubular Na^+^ flow past the macula densa, renal sympathetic nervous activity (RSNA), angiotensin II receptor type 1 (AT1R)-bound angiotensin II (Ang II) concentration, plasma [Ca^2+^], and plasma [1,25(OH)_2_D_3_] on renin secretion rate. All these terms are equal to 1 in the baseline condition. For definitions of *N*_*rs*_, *v*_*md*−*sod*_, *v*_*rsna*_,and *v*_*AT*1*R*_ refer to Ref. (29). The terms *v*_*Ca*_ and 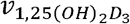 are defined below.

Although Ca^2^□ does not directly regulate renin secretion, its levels influence cAMP-stimulated renin secretion, with increased Ca^2^□ inhibiting and decreased Ca^2^□ enhancing this process. Acute hypercalcemia suppresses plasma renin activity (PRA) by acting on the calcium-sensing receptor (CaSR) (11). The JG cells, which express CaSR, reduce renin secretion when the receptor is activated (12). The basolateral surface of the JG cells is exposed to the renal cortical interstitium, so elevated Ca^2^□ levels in this area could directly stimulate the CaSR on JG cells, leading to a reduction in renin secretion. The effect of Ca^2+^ on renin secretion is given by

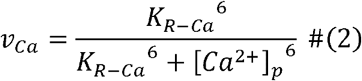

where *K*_*R*−*Ca*_ represents the sensitivity of renin secretion to plasma Ca^2+^ concentration ([*Ca*^2+^]_*p*_).

Renin secretion increases significantly in the presence of 1,25(OH)_2_D_3_ deficiency as well as toxicity. Zhou et al. reported a 1.5-fold increase in PRA in 1α-hydroxylase knockout mice compared to control (13). Treatment of 1α-hydroxylase knockout mice with 1,25(OH)_2_D_3_ led to the normalization of PRA (13). Thus, 1,25(OH)_2_D_3_ deficiency leads to upregulation of renin secretion. In addition, administration of high doses of calcitriol to male rats increased PRA by ∼2.5-fold (14). Thus, sufficiently high levels of 1,25(OH)_2_D_3_ also upregulates renin secretion.

We defined 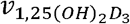 such that (i) when 1,25(OH)_2_D_3_ concentration is within the range of 80-170 pM, 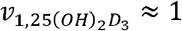, (ii) when 1,25(OH)_2_D_3_ concentration is less than 80 pM, 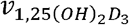gradually increases to a maximum of 1.5-fold as 1,25(OH)_2_D_3_ concentration approaches zero, and (iii) when 1,25(OH)_2_D_3_ concentration is greater than 170 pM, 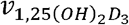 gradually increases to a maximum of 2-fold. Thus, the effect of 1,25(OH)_2_D_3_ on renin secretion is given by

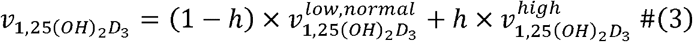

where 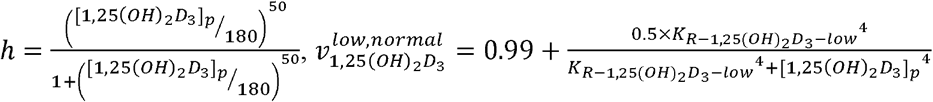, and 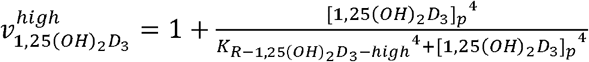. The parameters, 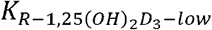 and 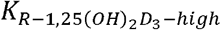, represent the sensitivity of renin secretion to low and high plasma 1,25(OH)_2_D_3_ concentrations ([1,25(*OH*_2_)*D*_3_]_*p*_), respectively.

### 2.2 Aldosterone secretion

Normalized aldosterone secretion (*N*_*als*_) is given by

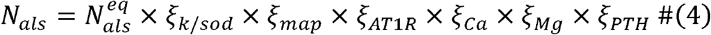

ALD secretion is affected by the K^+^ to Na^+^ ratio (*ξ*_*k*/*sod*_), mean arterial pressure (*ξ*_*map*_), AT1R-bound Ang II (*ξ*_*AT*1*R*_), Ca^2+^ (*ξ*_*Ca*_), Mg^2+^ (*ξ*_*Mg*_), and PTH (*ξ*_*PTH*_). All these terms are equal to 1 in the baseline condition. For definitions of 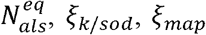, and *ξ*_*AT*1*R*_ refer to Ref. (29). The terms *ξ*_*Ca*_, *ξ*_*Mg*_, and *ξ*PTH are defined below.

The influence of extracellular Ca^2+^ on aldosterone synthesis and secretion was first reported in rat zona glomerulosa (ZG) cells (15). There were no changes in both cAMP and aldosterone levels when rat ZG cells were placed into Ca^2+^-deprived media. However, after rat ZG cells were treated with additional extracellular Ca^2+^, the concentrations of both aldosterone and cAMP were elevated in a dose-dependent manner. These findings suggested that extracellular Ca^2+^ level could serve as an independent stimulator of aldosterone secretion. We model the effect of Ca^2+^ on aldosterone secretion as

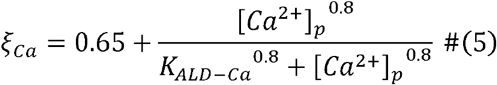

where *K*_*ALD*−*Ca*_ represents the sensitivity of aldosterone secretion to plasma Ca^2+^ concentration. AT1R-bound Ang II stimulates aldosterone secretion by increasing Ca^2+^ influx through the various voltage-gated Ca^2+^ channels. Mg^2+^ affects aldosterone secretion by blocking the voltage-gated Ca^2+^ channels which reduce Ang II-stimulated aldosterone secretion (30). In adrenal cells in culture, increased extracellular Mg^2+^ reduced Ang II-stimulated aldosterone release (16). This effect was also observed in vivo, as rats given an infusion of Ang II in combination with Mg^2+^ sulfate exhibited diminished plasma aldosterone, as compared with rats given Ang II alone (17). In addition, in a clinical study, participants placed on a very low Mg^2+^ diet for 3 weeks showed increased Ang II-stimulated aldosterone secretion (31). This increase in aldosterone secretion was attenuated by acute intravenous Mg^2+^ repletion. Together, these studies suggest that extracellular Mg^2+^ decreases sensitivity of adrenal glomerulosa cells to Ang II-stimulated aldosterone secretion. We model the effect of Mg^2+^ on aldosterone secretion as

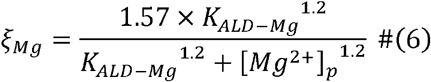

where *K*_*ALD*−*Mg*_ represents the sensitivity of aldosterone secretion to plasma Mg^2+^ concentration ([*Mg*^2+^] _*p*_).

The presence of type 1 PTH receptor has been demonstrated in both rat and human adrenal gland (18,19,32,33). PTH binds to PTH receptor and facilitates Ca^2+^ influx through the voltage-gated Ca^2+^ channels, thus increasing Ang II-stimulated aldosterone secretion. We model the effect of PTH on aldosterone secretion as

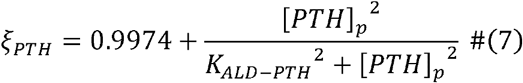

where *K*_*ALD*−*PTH*_ represents the sensitivity of aldosterone secretion to plasma PTH concentration ([_*PTH*_]_*p*_).

### 2.3 Effect of angiotensin II and aldosterone on PTH secretion

Human parathyroid cells have been found to express mineralocorticoid receptors (MRs) (23,33) and AT1Rs (23). The AT1R expression level was found to be ∼100-fold lower than the MR expression level (23). Lenzini et al. reported that at physiological Ca^2+^ concentrations cells exposed to aldosterone increased PTH secretion by 271% compared to control (23). This increase in PTH secretion was abolished on treatment with an MR blocker. In addition, at physiological Ca^2+^ concentrations cells exposed to Ang II increased PTH secretion by 267% and this increase was abolished on treatment with an AT1R antagonist (23). However, co-stimulation with both aldosterone and Ang II did not produce any additive increase in PTH secretion, with PTH secretion increasing by 225% compared to control (23).

Based on the observations reported in (23), we model the effect of aldosterone and Ang II on PTH secretion as

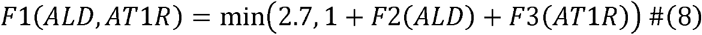

where

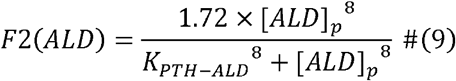

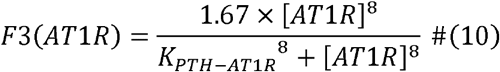

In Eqs. (9) and (10), [*ALD*] _*p*_ denotes plasma aldosterone concentration and [*AT*1*R*] denotes AT1R-bound Ang II concentration. *K*_*PTH* − *ALD*_ and *K*_*PTH*−*AT*1*R*_ denote the sensitivity of PTH secretion to aldosterone and AT1R-bound Ang II, respectively.

Equations (1) and (6) of the Ca^2+^-Mg^2+^ homeostasis model (28) representing the rates of change of parathyroid gland and plasma PTH concentrations, respectively, are modified as

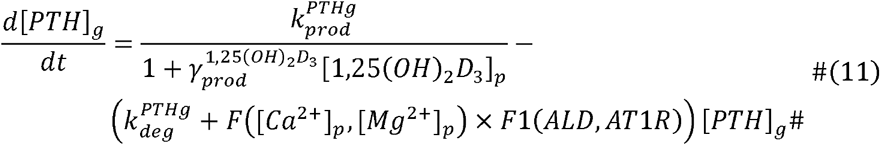

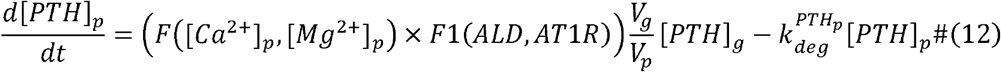

where [*PTH*]_*g*_ and [*PTH*]_*p*_ denote parathyroid gland and plasma PTH concentrations, respectively. For descriptions and values of all parameters in the above two equations refer to Table 2 of Ref. (28).

**Table 2.** Changes in model parameters to simulate five model instances of hypertension, each with different primary trigger(s), labelled *RSNA, Renin, ALD, AA*, and *Combined*. RSNA, renal sympathetic nervous activity; ALD, aldosterone; AA, afferent arterial.

|  | *RSNA* | *Renin* | *ALD* | *AA* | *Combined* |
| --- | --- | --- | --- | --- | --- |
| Baseline RSNA | $\times 1.54$ | $\times 1.0$ | $\times 1.0$ | $\times 1.0$ | $\times 1.28$ |
| Renin secretion rate | $\times 1.0$ | $\times 6.0$ | $\times 1.0$ | $\times 1.0$ | $\times 1.28$ |
| ALD secretion rate | $\times 1.0$ | $\times 1.0$ | $\times 30$ | $\times 1.0$ | $\times 1.28$ |
| Baseline AA resistance | $\times 1.0$ | $\times 1.0$ | $\times 1.0$ | $\times 2.9$ | $\times 1.28$ |

### 2.4 Effect of calcium and magnesium on RSNA

The regulation of RSNA is given by

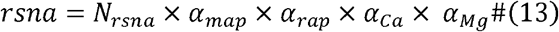

where *N*_*rsna*_ represents the normalized RSNA, and *α*_*map*_, *α*_*rap*_, *α*_*Ca*_, and *α*_*Mg*_ represent the effect of mean arterial pressure (MAP), right atrial pressure (RAP), Ca^2+^, and Mg^2+^ on RSNA.

All these terms are equal to 1 in the baseline condition. For definitions of *α*_*map*_ and *α*_*rap*_ refer to Ref. (29) and *α*_*Ca*_ and *α*_*Mg*_ are defined below.

Ca^2+^ influx through N-type Ca^2+^ channels in sympathetic nerves increases RSNA, which leads to increased release of norepinephrine (26). Mg^2+^ decreases RSNA and norepinephrine release by blocking N-type Ca^2+^ channels in sympathetic nerves (27). Thus, the balance between Ca^2+^ and Mg^2+^ is important for RSNA. We model the regulation of RSNA by Ca^2+^ and Mg^2+^ as

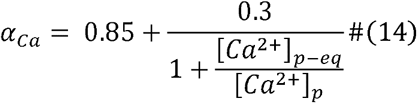

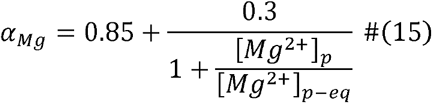

Where [*Ca*^2+^]_*p* −*eq*_ and [*Mg*^2+^]_*p* −*eq*_ denote the concentrations of plasma Ca^2+^ and Mg^2+^ at equilibrium, respectively.

### 2.5 Effect of angiotensin II on bone resorption

Bone resorption rate is defined as

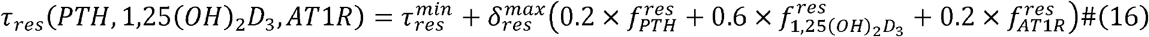

where 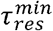 and 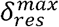 denote the minimal and maximal resorption rates, respectively, and their values can be found in Table 2 of Ref (28). The terms 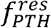 and 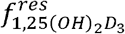 represent the effect of PTH and 1,25(OH)_2_D_3_ on resorption, respectively, and their definitions are provided in Ref (28). The term 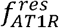 represents the effect of AT1R-bound Ang II on bone resorption and is defined below.

Bones express AT1R and Ang II binds to AT1R to activate osteoclasts through RANKL induction and hence promotes osteoporosis (34). Treatment with AT1R blocker ameliorates osteoporosis (34). We assume the following sigmoidal relation between bone resorption and AT1R-bound Ang II.

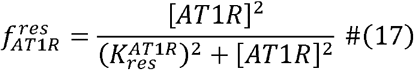

where 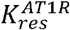 denotes the sensitivity of bone resorption to AT1R-bound Ang II.

### 2.6 Sodium reabsorption in the proximal tubule and thick ascending limb

Fractional reabsorption of Na^+^ in the proximal tubule and thick ascending limb is given by

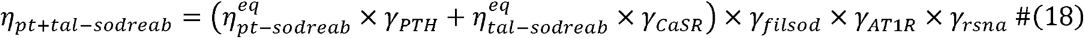

where 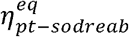 and 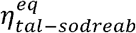 represent fractional Na^+^ reabsorption in the proximal tubule and thick ascending limb at equilibrium and are assumed to be 0.60 and 0.20, respectively. *γ*_*PTH*_, *γ*_*CaSR*_, *γ*_*filsod*_, *γ*_*AT*1*R*_, and *γ*_*rsna*_ denote the effect of PTH, CaSR, filtered Na^+^ load, AT1R-bound Ang II, and RSNA, respectively, on fractional Na^+^ reabsorption and are equal to 1 at baseline. The definitions of *γ*_*filsod*_, *γ*_*AT*1*R*_, and *γ*_*rsna*_ can be found in Ref. (29) and *γ*_*PTH*_ and *γ*_*CaSR*_ are defined below.

PTH inhibits Na^+^/H^+^ exchanger 3 (NHE3) in the proximal tubule and hence reduces proximal tubule Na^+^ reabsorption (24). We model the effect of PTH on fractional Na^+^ reabsorption in the proximal tubule as

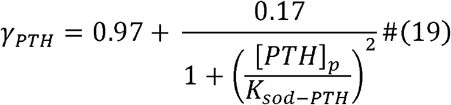

where *K*_*sod*−*PTH*_ represents the sensitivity of proximal tubule Na^+^ reabsorption to plasma [PTH]. Ca^2+^ inhibits Na^+^–K^+^–Cl^−^ cotransporter 2 (NKCC2) in the thick ascending limb through the CaSR and hence reduces Na^+^ reabsorption along this segment (25). We model the effect of Ca^2+^ on fractional Na^+^ reabsorption in the thick ascending limb as

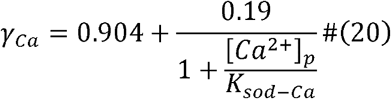

where *K*_*sod*−*Ca*_ represents the sensitivity of Na^+^ reabsorption in the thick ascending limb to plasma [Ca^2+^].

### 2.7 Effect of angiotensin II on arterial resistance

In vascular smooth muscle cells, Ang II binds to AT1R and increases intracellular Ca^2+^ both through Ca^2+^ influx from L-type Ca^2+^ channels and Ca^2+^ release from intracellular stores. Administration of angiotensin converting enzyme (ACE) inhibitor to spontaneously hypertensive rats reduced peripheral resistance by ∼40% (35) and Ang II infusion to male Wistar rats increased peripheral resistance by ∼140% (36). Based on these observations, we model the effect of AT1R-bound Ang II on arterial resistance as

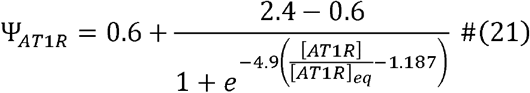

where [*AT*1*R*]_*eq*_ denotes the concentration of AT1R-bound Ang II at equilibrium.

### 2.8 Effect of calcium, magnesium, and parathyroid hormone on vascular resistance

Increased Ca^2+^ influx into vascular smooth muscle cells increases vascular resistance. Low and high Ca^2+^ supplementation in male Sprague-Dawley rats decreased and increased vascular resistance by 13.5% and 7%, respectively (20). Accordingly, we model the effect of plasma Ca^2+^ on vascular resistance as

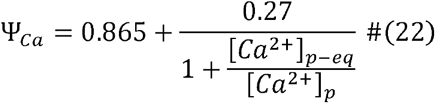

where [*Ca*^2+^]_*p* −*eq*_ denotes the concentration of plasma Ca^2+^ at equilibrium.

Mg^2+^ causes vasodilation of vascular smooth muscle cells by blocking Ca^2+^ influx through L-type Ca^2+^ channels. High extracellular Mg^2+^ reduced myogenic tone in wild-type male mice (21). We model the effect of plasma Mg^2+^ on vascular resistance as

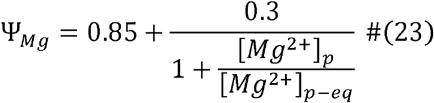

where [*Mg*^2+^]_*p* −*eq*_ denotes the concentration of plasma Ca^2+^ at equilibrium.

PTH binds to PTHr-1 and increases Ca^2+^ influx into vascular smooth muscle cells (22). We modeled the effect of PTH on vascular resistance by assuming that at very low PTH vascular resistance would decrease by 20% and at high PTH vascular resistance would increase by 20%.

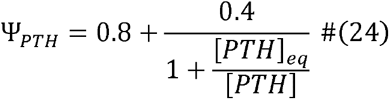

where [*PTH*]_*eq*_ denotes the concentration of plasma PTH at equilibrium.

The regulation of arterial resistance is given by

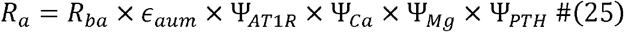

where *R*_*ba*_ denotes the basic arterial resistance and *ϵ*_*aum*_ denotes the autonomic multiplier effect and their definitions can be found in Ref. (29). The parameters Ψ_*AT*1*R*_, Ψ_*Ca*_, Ψ_*Mg*_, and Ψ_*PTH*_ denote the effect of [AT1R-bound Ang II], [Ca^2+^], [Mg^2+^], and [PTH] on vascular resistance and are defined in Eqs. (21), (22), (23), and (24), respectively.

The regulation of afferent arteriolar resistance is defined as

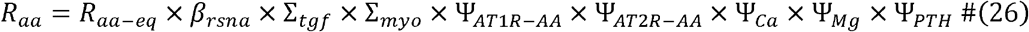

where *R*_*aa*− *eq*_ denotes the afferent arteriolar resistance at equilibrium. The parameters *β*_*rsna*_, Σ_*tgf*_, Σ_*myo*_, Σ_*AT*1*R−AA*_, Ψ_*AT*2*R*−*AA*_, and ΨATR-AA denote the effect of RSNA, tubuloglomerular feedback (TGF) signal, myogenic response, [AT1R-bound Ang II], and [AT2R-bound Ang II] on afferent arteriolar resistance, respectively, and their definitions can be found in Ref. (29). The parameters Ψ_*Ca*_, Ψ_*Mg*_, and Ψ_*PTH*_ denote the effect of [Ca^2+^], [Mg^2+^], and [PTH] on afferent arteriolar resistance and are defined in Eqs. (22), (23), and (24), respectively.

### 2.9 Simulating hypertension

Hypertension is a multifactorial disease that may involve a variety of triggers, including overactive RSNA, RAAS, or arterial stiffening. The model parameters adjusted to simulate hypertensive stimuli are the equilibrium values for RSNA, renin secretion rate, aldosterone secretion rate, and afferent arteriole resistance. We consider five hypertensive cases, featuring primarily an overactive RSNA, increased renin secretion, increased aldosterone secretion, increased vascular tone, or a combination of these stimuli. These models are referred to as HTN-RSNA, HTN-Renin, HTN-ALD, HTN-AA, and HTN-Combined, respectively. For each hypertensive case, parameters were adjusted so that the MAP predicted for the hypertensive model is approximately 120 mmHg. These parameter sets are shown in Table 2.

## 3. Results

### 3.1 Sensitivity analysis

We performed a local sensitivity analysis by varying each model parameter by ±5% and computing the corresponding steady state. The percentage changes in RSNA, glomerular filtration rate (GFR), MAP, aldosterone concentration, PRA, [PTH], [1,25(OH)_2_D_3_], [Mg^2+^], and [Ca^2+^] corresponding to 5% increase in parameter values are shown in Fig. 2.

**Figure 2:**
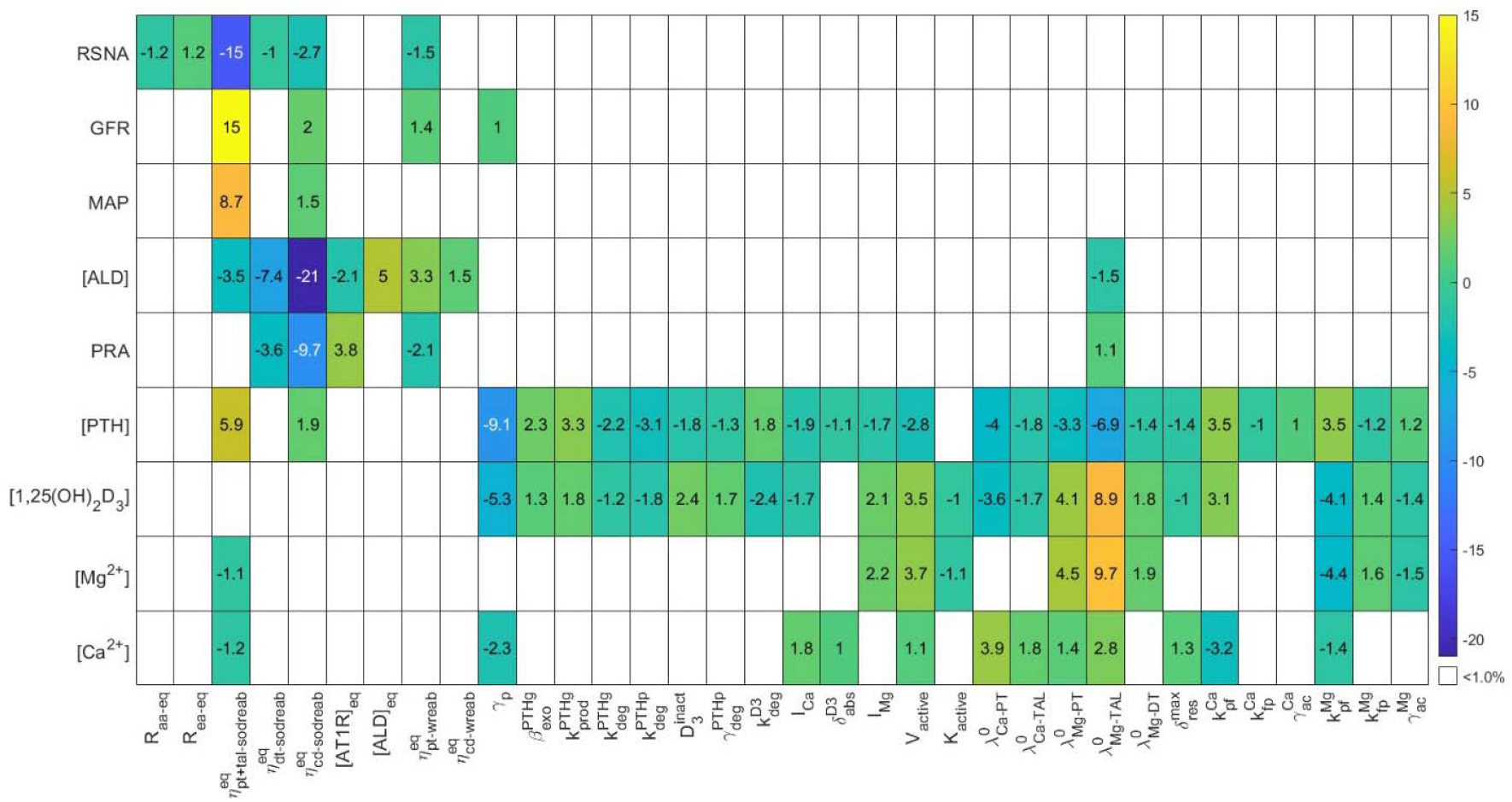
Local sensitivity analysis conducted by increasing individual parameters by 5%. The resulting percent change in model steady state concentrations from baseline is presented here. White indicates the resulting change was less than 1%. RSNA, renal sympathetic nervous activity; GFR, glomerular filtration rate; MAP, mean arterial pressure; [ALD], plasma aldosterone concentration; PRA, plasma renin activity; [PTH], plasma PTH concentration; [1,25(OH)_2_D_3_], plasma 1,25(OH)_2_D_3_ concentration; [Mg^2+^], plasma Mg^2+^ concentration; [Ca^2+^], plasma Ca^2+^ concentration.

From Fig. 2 we see that the parameters, 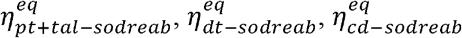, and 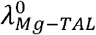, have the largest impacts on different model variables. A 5% increase in the fractional reabsorption rates of Na^+^ in the proximal tubule and thick ascending limb 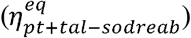, distal tubule 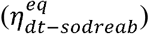, and collecting duct 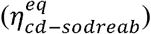 causes significant change in aldosterone concentration. Additionally, 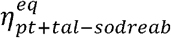 has significant impact on GFR (since it determines the macula densa Na^+^ flow), RSNA, and MAP (by regulating water reabsorption and consequently extracellular fluid volume). A 5% increase in minimal thick ascending limb fractional Mg^2+^ reabsorption 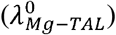 causes a significant change in plasma [PTH], [1,25(OH)_2_D_3_], [Mg^2+^], and [Ca^2+^]. For instance, the increase in 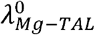 significantly elevates plasma [Mg^2+^] and [Ca^2+^], while suppressing plasma [PTH]. This occurs because increased plasma [Mg^2+^] directly stimulates 1,25(OH)_2_D_3_ production, which in turn enhances intestinal Ca^2+^ absorption, thereby increasing plasma [Ca^2+^]. Consequently, [PTH] decreases due to inhibition from both increased [1,25(OH)_2_D_3_] and elevated [Mg^2+^] and [Ca^2+^]. In contrast, a similar change in fractional Ca^2+^ reabsorption in the proximal tubule 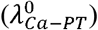 has minimal impact on these variables due to a negative feedback loop: increased plasma [Ca^2+^] inhibits PTH and 1,25(OH)_2_D_3_, dampening further Ca^2+^ changes. Unlike Ca^2+^, the feedback loop of Mg^2+^ with 1,25(OH)_2_D_3_ is reinforcing, amplifying the impact of renal Mg^2+^ reabsorption changes. Local sensitivity analysis conducted by decreasing individual parameters by 5% also showed similar trends.

### 3.2 Effect of different hypertensive stimuli

As described above, we adjusted selected parameters to investigate the models’ responses to different hypertensive stimuli: overactive RSNA, increased renin secretion, increased aldosterone secretion, increased vascular tone, or a combination of these stimuli (Table 2). Also, these parameters were chosen to yield MAP of approximately 120 mmHg, corresponding to an increase of 16% (Fig. 3A). Model predictions for each of these stimuli are shown in Fig. 3.

**Figure 3:**
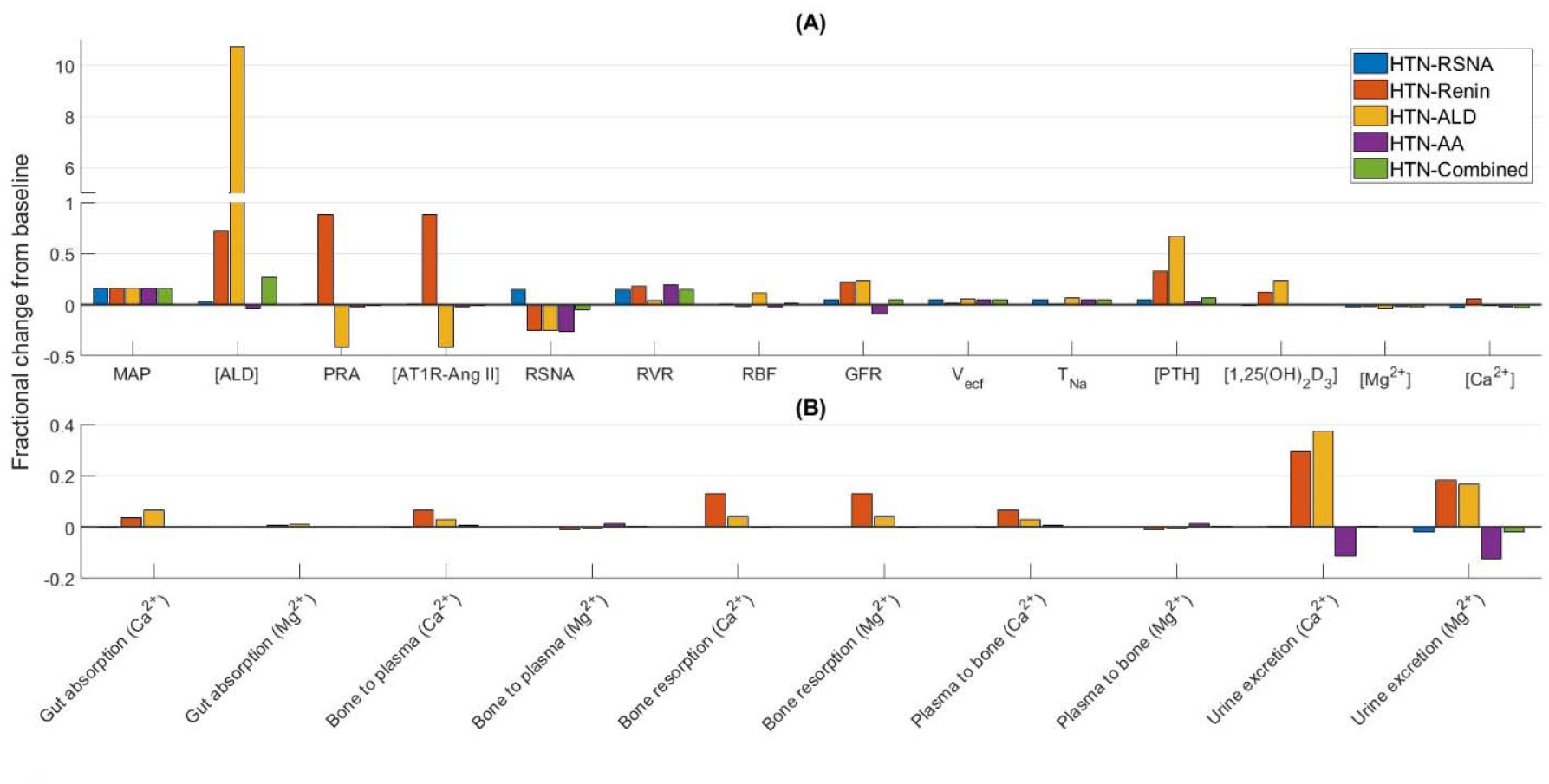
Fractional change from baseline (denoted by grey line at zero) of model variables (A) and Ca^2+^-Mg^2+^ fluxes (B) under five hypertensive stimuli: overactive RSNA (HTN-RSNA), increased renin secretion (HTN-Renin), increased aldosterone secretion (HTN-ALD), increased vascular tone (HTN-AA), and a combination of these stimuli (HTN-Combined). Bone to plasma and plasma to bone fluxes represent the exchange of Ca^2+^ and Mg^2+^ between plasma and the fast bone pool; bone resorption represents the release of Ca^2+^ and Mg^2+^ from the slow bone pool into plasma. MAP, mean arterial pressure; [ALD], aldosterone concentration; PRA, plasma renin activity; [AT1R-Ang II], plasma AT1R-bound Ang II concentration; RSNA, renal sympathetic nervous activity; RVR; renal vascular resistance; RBF, renal blood flow; GFR, glomerular filtration rate; V_ecf_, extracellular fluid volume; T_Na_, total plasma sodium; [PTH], plasma PTH concentration; [1,25(OH)_2_D_3_], plasma 1,25(OH)_2_D_3_ concentration; [Mg^2+^], plasma Mg^2+^ concentration; [Ca^2+^], plasma Ca^2+^ concentration.

In the HTN-RSNA case, RSNA induces vasoconstriction and directly stimulates proximal tubule Na^+^ reabsorption. Taken in isolation, afferent arteriolar constriction would lower GFR. However, the resulting reduced Na^+^ flow at the macula densa inhibits the TGF signal and causes the afferent arteriole to slightly dilate. The higher MAP would also increase GFR. Together, these factors yield a GFR that is slightly higher than the normotensive values (Fig. 3(A)). RSNA-induced hypertension does not cause any significant changes in plasma Ca^2+^, Mg^2+^, PTH, and 1,25(OH)_2_D_3_ levels or on any of Ca^2+^-Mg^2+^ fluxes.

In the HTN-Renin case (hyperreninemia), increased renin secretion elevates the level of AT1R-bound Ang II, which constricts both the afferent and efferent arterioles, but preferentially the latter. As a result, while renal blood flow was predicted to be almost unchanged from baseline, GFR was notably higher (22%; Fig. 3(A)). The significantly increased aldosterone stimulates PTH secretion (39,40), thereby increasing plasma [PTH] by 33%. On one hand, this increased [PTH] inhibits proximal tubular Ca^2+^ and Mg^2+^ reabsorption, while on the other hand it increases Ca^2+^ and Mg^2+^ reabsorption along the thick ascending limb and distal tubule. Now, majority of Ca^2+^ reabsorption occurs along the proximal tubule, while majority of Mg^2+^ reabsorption occurs along the thick ascending limb. Hence, combined with the higher filtered load, our model predicts a notably higher increase in urinary Ca^2+^ excretion than in urinary Mg^2+^ excretion (Fig. 3(B)). Further, bone resorption increases by 13% under PTH, calcitriol, and AT1R-bound Ang II stimulation. Together, these factors do not cause any significant change in plasma Mg^2+^ and Ca^2+^ concentrations.

The HTN-ALD case simulates primary hyperaldosteronism, where the renin secretion rate remains normal, but there is hypersecretion of aldosterone, with a 30-fold increase above baseline.

Aldosterone secretion rate in primary aldosteronism is difficult to measure. However, we calculated the ratio of plasma aldosterone concentration (PAC) (ng/dL) to plasma renin activity (PRA) (pmol/L/min), which is commonly used in the diagnosis of primary aldosteronism. The proposed cut-off values for PAC/PRA ratio in the literature are 1.6, 2.5, and 3.1 ng/dL per pmol/L/min (41). We obtained a PAC/PRA ratio of 1.86 ng/dL per pmol/L/min with the predicted plasma aldosterone concentration and PRA values. The high levels of aldosterone cause the kidneys to retain Na^+^ and increase blood volume, which in turn signals the kidneys to decrease renin production. Our model predicts a 41% decrease in plasma renin activity and AT1R-bound Ang II (Fig. 3(A)). The high aldosterone level also causes a 67% increase in PTH concentration (Fig. 3(A)) (39,40). This in turn inhibits proximal tubular Ca^2+^ and Mg^2+^ reabsorption, while increasing their reabsorption along the thick ascending limb. Consequently, our model predicts a 37% increase in urinary Ca^2+^ excretion (40,42) while urinary Mg^2+^ excretion is predicted to increase by 17% (43) (Fig. 3(B)). Under the combined effect of increased PTH (67%), increased calcitriol (23%), and decreased AT1R-bound Ang II (41%), bone resorption increases by 3.8% (39,44,45) (Fig 3(B)). Together, these factors keep plasma Mg^2+^ and Ca^2+^ concentrations near the baseline values (40,43).

Afferent arteriole constriction in the HTN-AA case lowers GFR, whereas the increased MAP increases GFR. Under the influence of these two opposing factors GFR decreases by 8.5%. This in turn reduces proximal tubule Na^+^ and water reabsorption by a small amount. The increased Na^+^ delivery to the downstream segments increases Na^+^ reabsorption in the distal tubule and collecting duct. Hence, total plasma Na^+^ and extracellular fluid volume increase slightly. In addition, the reduced GFR lowers urinary Ca^2+^ and Mg^2+^ excretions.

The HTN-Combined case involves an overactive RSNA, an overactive RAAS, and increased vascular tone, with the strength of these stimuli chosen so that the predicted MAP is ∼120 mmHg (Table 2). The high aldosterone level increases [PTH] by 6.4%. However, it does not cause any notable change in any of the Ca^2+^ and Mg^2+^ fluxes.

Thus, the HTN-Renin, HTN-ALD, and HTN-AA cases have significant effect on Ca^2+^ and Mg^2+^ fluxes. However, plasma Ca^2+^ and Mg^2+^ concentrations are predicted to remain within their respective physiological ranges for all hypertensive stimuli.

### 3.3 Effect of Mg^2+^, Ca^2+^, and vitamin D_3_ deficiency

Next, we simulated 70% dietary Mg^2+^ intake (*I*_*Mg*_) restriction, 70% dietary Ca^2+^ intake (*I*_*Ca*_) restriction, and 70% 25(OH)D (inactive form of vitamin D_3_) deficiency for 1 month. Model predictions are shown in Fig. 4.

**Figure 4:**
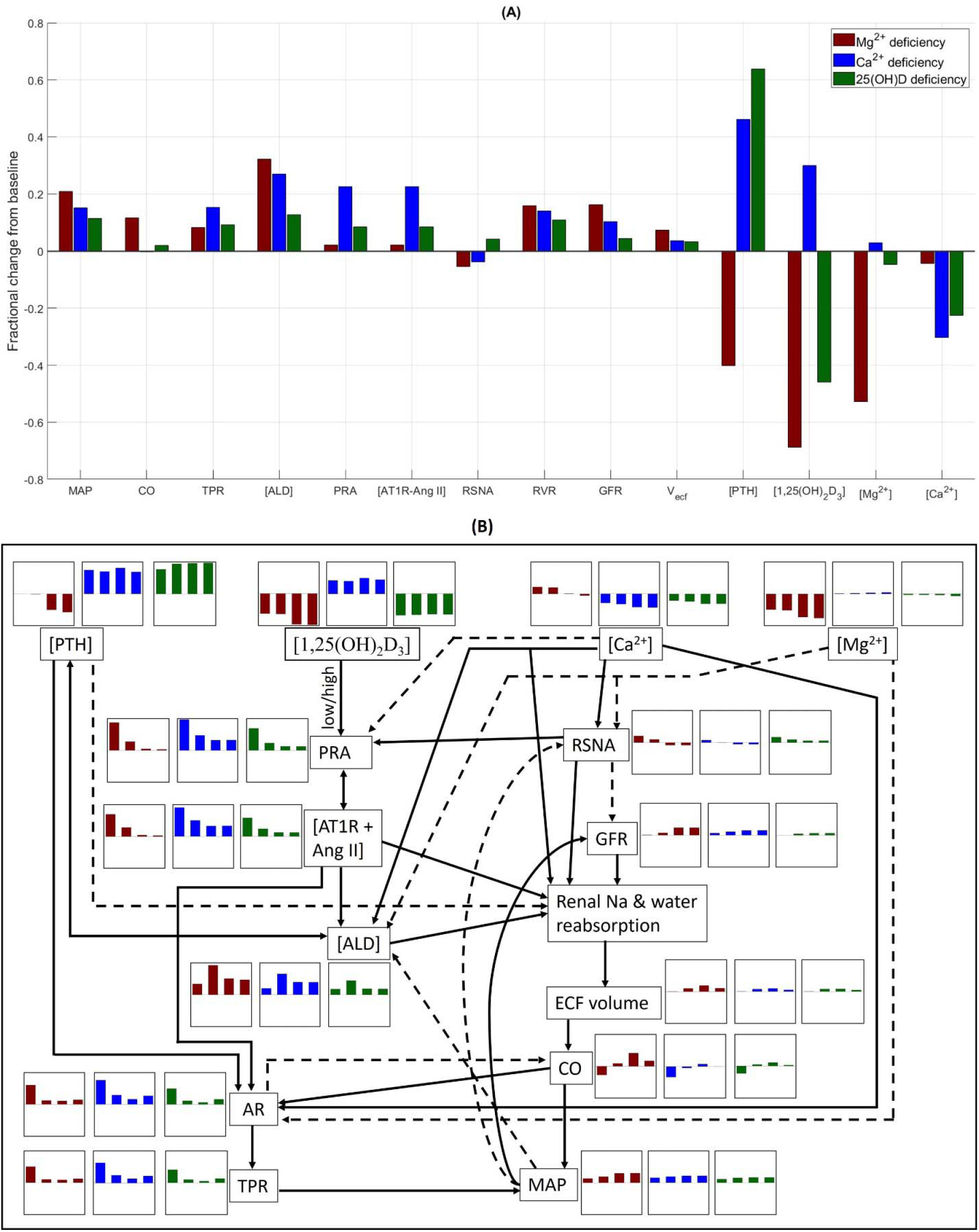
(A) Fractional change from baseline (denoted by grey line at zero) of model variables under 70% dietary Mg^2+^ intake (*I*_*Mg*_) restriction, 70% dietary Ca^2+^ intake (*I*_*Ca*_) restriction, and 70% 25(OH)D (inactive form of vitamin D_3_) deficiency for 1 month. (B) Interplay between key variables that are affected by dietary Mg^2+^, dietary Ca^2+^, and 25(OH)D deficiency. Each variable is accompanied by their change over time for each of the three cases. Maroon bar plots, dietary Mg^2+^ deficiency; blue bar plots, dietary Ca^2+^ deficiency; green bar plots, 25(OH)D deficiency. The Y-axis range is [-0.7, 0.7]. MAP, mean arterial pressure; CO, cardiac output; TPR, total peripheral resistance; [ALD], aldosterone concentration; PRA, plasma renin activity; [AT1R-Ang II], AT1R-bound Ang II concentration; RSNA, renal sympathetic nervous activity; RVR; renal vascular resistance; GFR, glomerular filtration rate; V_ecf_, extracellular fluid volume; [PTH], plasma PTH concentration; [1,25(OH)_2_D_3_], plasma 1,25(OH)_2_D_3_ concentration; [Mg^2+^], plasma Mg^2+^ concentration; [Ca^2+^], plasma Ca^2+^ concentration; AR, arterial resistance.

Our model predicts a 21% increase in MAP from 103 mmHg to 125 mmHg following 1 month of dietary Mg^2+^ restriction (Fig. 4(A)). By contrast, 1 month of dietary Ca^2+^ restriction and 25(OH)D deficiency have comparatively lesser effect on MAP (increases to 119 mmHg and 114 mmHg, respectively). Now, why does Mg^2+^ deficiency have a higher effect on MAP compared to Ca^2+^ and 25(OH)D deficiency? Figure 4(B) shows the key variables and the interplay between them to answer the above question. Each variable is accompanied by their change over time for each of the three cases.

Now, MAP is modeled as the product of cardiac output and total peripheral resistance; the relative change in these two factors determines the change in MAP. Dietary Mg^2+^ deficiency markedly lowers plasma [Mg^2+^], which in turn inhibits PTH secretion. Hence plasma [PTH] drops by 40%. This in turn partially removes the inhibitory effect of PTH on proximal tubule Na^+^ reabsorption. From Fig 4(B), we see that RSNA (maroon bar plot) initially rises under the effect of lower plasma [Mg^2+^]. However, as MAP rises, its inhibitory effect brings down RSNA. Also, GFR (Fig. 4(B), maroon bar plot) does not change initially, but as MAP rises it increases renal blood flow and hence GFR. Thus, proximal tubule Na^+^ and water reabsorption is enhanced under the combined effect of increased GFR and removal of the inhibitory effect of PTH. Additionally, the lowered plasma [Mg^2+^] removes the inhibitory effect on aldosterone secretion causing an increase in plasma aldosterone levels. The high aldosterone levels enhance Na^+^ and water reabsorption in the distal tubule and collecting duct. Consequently, the extracellular fluid volume increases, which subsequently increases cardiac output by 12%. Now, the increased cardiac output increases the arterial resistance. This is because when more blood flows through any tissue of the body than is required by that tissue for its specific function, the local resistance to blood flow increases progressively to bring the blood flow back towards normal. On the other hand, the lower [PTH] decreases arterial resistance and lower [Mg^2+^] increases arterial resistance. Under the combined effect of these and cardiac output, arterial resistance increases which in turn increases the total peripheral resistance by 8.3%. Thus, dietary Mg^2+^ deficiency increases cardiac output and total peripheral resistance, which together cause a 21% increase in MAP. In fact, several observational studies, clinical trials, and meta-analyses have shown an inverse relationship between dietary Mg^2+^ intake and hypertension (46–50).

By contrast, dietary Ca^2+^ deficiency significantly increases plasma [PTH] due to the drop in plasma [Ca^2+^]. This reinforces the inhibitory effect of PTH on proximal tubule Na^+^ reabsorption. In addition, RSNA (Fig. 4(B), blue bar plot) decreases under the combined effect of lower plasma [Ca^2+^] and rising MAP. The increasing MAP also increases GFR (Fig. 4(B), blue bar plot). Furthermore, the lower plasma [Ca^2+^] removes the inhibitory effect on renin secretion; hence PRA increases, which in turn increases [AT1R-bound Ang II] (Fig. 4(B), blue bar plot). Together these factors slightly increase proximal tubule Na^+^ and water reabsorption. Additionally, the increased aldosterone (under the effect of increased [PTH] and [AT1R-bound Ang II]) increases Na^+^ and water reabsorption in the distal tubule and collecting duct. Hence, the extracellular fluid volume increases, which in turn increases cardiac output. On the other hand, the higher [AT1R-bound Ang II] and [PTH] increase arterial resistance and the lower [Ca^2+^] decreases arterial resistance. Together these factors increase total peripheral resistance by 15%. This increased resistance inhibits cardiac output. Consequently, cardiac output does not change from the baseline value. Thus, during dietary Ca^2+^ deficiency the 15% increase in MAP is primarily due to increased total peripheral resistance. Several observational studies, clinical trials, and meta-analyses have shown an inverse relationship between dietary Ca^2+^ intake and hypertension (51–53).

Calcitriol (1,25(OH)_2_D_3_) deficiency following 25(OH)D restriction lowers intestinal absorption of Ca^2+^ and hence causes a drop in plasma [Ca^2+^]. This in turn increases PTH secretion. The decreased plasma [1,25(OH)_2_D_3_] also partially removes the inhibitory effect on PTH production. Hence, we see a higher rise in plasma [PTH] compared to the dietary Ca^2+^ deficiency case. In calcitriol deficiency, RAAS, renal Na^+^ and water reabsorption, and peripheral resistance undergo similar changes as in dietary Ca^2+^ deficiency, though the impact is lower. Consequently, MAP increases by only 11% (54–56).

Thus, according to our model predictions the factor that causes Mg^2+^ deficiency to have a higher effect on MAP compared to Ca^2+^ and 25(OH)D deficiency is the increased cardiac output. During Mg^2+^ deficiency, changes in GFR, RSNA, [PTH], [AT1R-bound Ang II], and aldosterone significantly increase renal Na^+^ and water reabsorption, leading to a considerable increase in extracellular fluid volume and thus cardiac output. On the other hand, during Ca^2+^ and 25(OH)D deficiency, changes in these factors cause modest changes in extracellular fluid volume and hence cardiac output does not change significantly from baseline.

### 3.4 Primary hyperparathyroidism

Primary hyperparathyroidism was simulated by increasing the baseline PTH synthesis rate 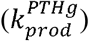 by factors of 2, 3, 5, 7, 10. Shown in Fig. 5 are the predicted steady-state values of key model variables and fluxes.

**Figure 5:**
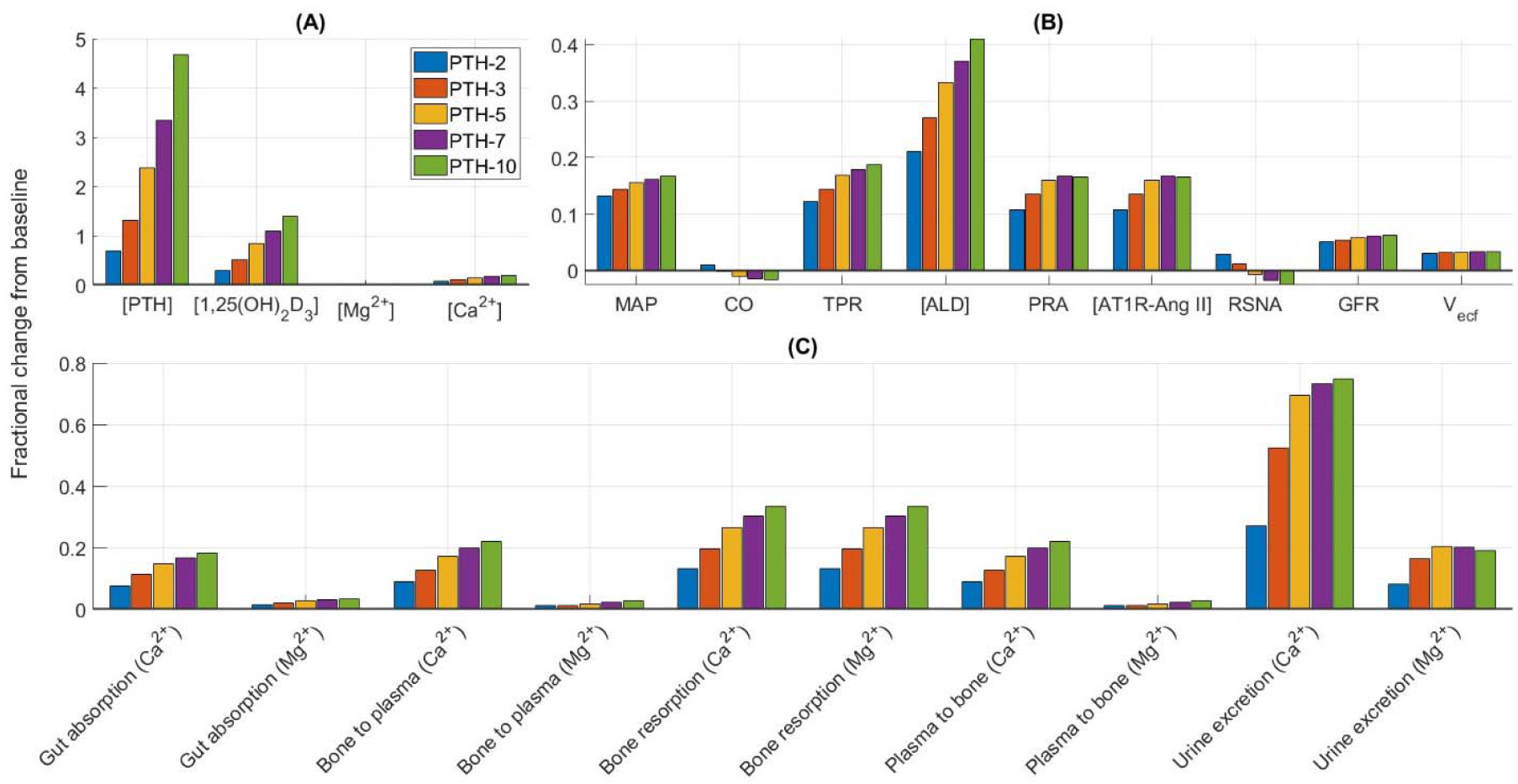
Fractional change from baseline (denoted by grey line at zero) of model variables (A, B) and Ca^2+^-Mg^2+^ fluxes (C) after increasing the baseline PTH synthesis rate 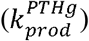 by factors of 2, 3, 5, 7, 10. [PTH], plasma PTH concentration; [1,25(OH)_2_D_3_], plasma 1,25(OH)_2_D_3_ concentration; [Mg^2+^], plasma Mg^2+^ concentration; [Ca^2+^], plasma Ca^2+^ concentration; MAP, mean arterial pressure; CO, cardiac output; TPR, total peripheral resistance; [ALD], aldosterone concentration; PRA, plasma renin activity; [AT1R-Ang II], AT1R-bound Ang II concentration; RSNA, renal sympathetic nervous activity; GFR, glomerular filtration rate; V_ecf_, extracellular fluid volume.

Our model predicts plasma [PTH] to increase by 70%, 130%, 237%, 334%, and 468%, respectively, following 2-, 3-, 5-, 7-, and 10-fold increase in 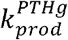. Since PTH increases the synthesis of 1,25(OH)_2_D_3_ we see a significant increase in plasma [1,25(OH)_2_D_3_]. This in turn greatly enhances intestinal Ca^2+^ absorption, while intestinal Mg^2+^ absorption increases slightly. This is because 1,25(OH)_2_D_3_ regulates 45% of intestinal Ca^2+^ absorption but only 12% of intestinal Mg^2+^ absorption. Further, bone resorption increases under the combined stimulation of PTH and 1,25(OH)_2_D_3_. Now, PTH inhibits proximal tubular Ca^2+^ and Mg^2+^ reabsorption, while stimulating Ca^2+^ and Mg^2+^ reabsorption along the thick ascending limb and distal tubule. Now, majority of Ca^2+^ reabsorption occurs along the proximal tubule, while majority of Mg^2+^ reabsorption occurs along the thick ascending limb. Hence, combined with the higher filtered load, our model predicts a notably higher increase in urinary Ca^2+^ excretion than in urinary Mg^2+^ excretion. Together these factors cause plasma [Ca^2+^] to increase by 7% (1.32 mM), 11% (1.37 mM), 15% (1.41 mM), 18% (1.45 mM), and 20% (1.48 mM), respectively, for 2-, 3-, 5-, 7-, and 10-fold increase in 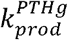. Thus, in all five cases, plasma [Ca^2+^] is above its physiological range (1.1-1.3 mM) indicating hypercalcemia (57). By contrast, the model predicts no change in plasma [Mg^2+^] (58).

Now let us analyze the effect on RAAS. The elevated plasma [Ca^2+^] inhibits renin secretion, whereas the elevated plasma [1,25(OH)_2_D_3_] increases renin secretion. Under their combined effect PRA increases significantly, which in turn increases AT1R-bound Ang II and aldosterone. The increased PTH and Ca^2+^ also increase aldosterone secretion. Thus, plasma aldosterone increases by 21%, 27%, 34%, 37%, and 42%, respectively, for 2-, 3-, 5-, 7-, and 10-fold increase in 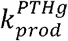 (33). Renal Na^+^ and water reabsorption increases under the stimulation of AT1R-bound Ang II and aldosterone. An important point to note is that though PTH indirectly enhances Na^+^ reabsorption through the RAAS, it also directly inhibits proximal tubule Na^+^ reabsorption. Because of these two opposing factors, extracellular fluid volume and total plasma Na^+^ increase only slightly. Additionally, the higher [AT1R-bound Ang II], [PTH], and [Ca^2+^] significantly increase arterial resistance and consequently total peripheral resistance. Thus, the increased extracellular fluid volume increases cardiac output, whereas the increased arterial resistance inhibits cardiac output and together these two factors keep cardiac output close to the baseline value. Consequently, MAP increases by 13% (116 mmHg), 14% (117 mmHg), 15.5% (119 mmHg), 16% (119.5 mmHg), and 17% (121 mmHg), respectively, for 2-, 3-, 5-, 7-, and 10-fold increase in 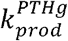. Thus, primary hyperparathyroidism causes hyperaldosteronism (33,59) and increases the risk of hypertension. In fact primary hyperparathyroidism has been associated with increased risk of hypertension, with a prevalence ranging from 40-65% and parathyroidectomy has resulted in substantial fall in both mean systolic and diastolic blood pressures (60–64).

Osteopenia and osteoporosis are also frequent complications of primary hyperparathyroidism (65–67). Our model predicts significant increase in bone resorption due to increased PTH, calcitriol, and AT1R-bound Ang II.

## 4 Discussion

We have developed an integrated model of blood pressure regulation and Ca^2^□-Mg^2^□ homeostasis to gain insights into the complex interactions that govern these physiological processes. Our study highlights the bidirectional relationship between these two systems, where alterations in one can lead to significant changes in the other. Specifically, we demonstrated how dysregulation in Ca^2+^ and Mg^2+^ levels can affect blood pressure control, and how dysregulation in blood pressure control, such as in hypertension, can impact Ca^2^□ and Mg^2^□ homeostasis.

Hypertension has been associated with elevated plasma PTH (68), increased Ca^2+^ excretion (68,69), and reduced bone mineral density (70). We used our model to simulate hypertension using different stimuli, namely, increased RSNA, increased renin secretion, increased aldosterone secretion, increased vascular tone, and a combination of all these stimuli. The model simulations, as depicted in Fig. 3, suggest that hypertensive states driven by elevated renin (HTN-Renin) and aldosterone (HTN-ALD) secretions significantly influence PTH, calcitriol, and the renal handling of Ca^2+^ and Mg^2+^. Notably, both conditions exhibit increased urinary excretion of Ca^2+^ and Mg^2+^, as well as elevated bone resorption (Fig. 3). Primary hyperaldosteronism, which results from increased aldosterone secretion, has been associated with hyperparathyroidism (39,40) and increased urinary excretions of Ca^2+^ (40,42) and Mg^2+^ (43), although plasma Ca^2+^ and Mg^2+^ remain normal (40,43). In fact, primary aldosteronism has been associated with nephrocalcinosis (71) and nephrolithiasis (72) caused by hypercalciuria. In addition, primary aldosteronism also causes bone loss and reduced bone mineral density (39,44,45). Now, hyperreninemia, caused by increased renin secretion, has been associated with hyperaldosteronism (73,74). Thus, the elevated aldosterone levels in hyperreninemia also increase PTH levels and hence urinary excretions of Ca^2+^ and Mg^2+^. Our model predicts a significantly higher increase in bone resorption rate in the HTN-Renin case compared to the HTN-ALD case (Fig. 3). This is because increased renin secretion increases ATIR-bound Ang II, which increases the risk of osteoporosis. In fact, the use of angiotensin-converting enzyme (ACE) inhibitors have been associated with higher bone mineral density and reduced risk of fracture (75,76). Despite these changes in urinary excretion and bone resorption rate, the model simulations indicate that plasma Ca^2+^ and Mg^2+^ concentrations remain relatively stable across all simulated hypertensive conditions (HTN-RSNA, HTN-Renin, HTN-ALD, HTN-AA, HTN-Combined) (Fig. 3), suggesting that the body’s compensatory mechanisms effectively maintain mineral homeostasis even under pathological conditions.

Cross-sectional and longitudinal epidemiological studies have consistently reported an inverse relationship between dietary Mg^2+^ and blood pressure and/or hypertension (77–84). Additionally, two large meta-analyses of randomized trials reported that Mg^2+^ supplementation significantly lowers blood pressure (48,49). Hypomagnesemia has also been associated with pre-eclampsia (85–87), a pregnancy specific hypertensive disorder, and Mg^2+^ supplementation has been reported to reduce the risk of eclampsia in pregnant women by over 50% (88). Thus, all these studies highlight the importance of Mg^2+^ homeostasis in blood pressure regulation. With our model simulations, we investigated the mechanisms responsible for increased mean arterial pressure during Mg^2+^ deficiency. Our model simulations showed that Mg^2+^ deficiency increased both cardiac output and total peripheral resistance (Fig. 4). The increase in cardiac output was primarily due to decreased PTH and increased aldosterone, which increased renal Na^+^ and water reabsorption. Total peripheral resistance increased mainly due to increased cardiac output and removal of the inhibitory effect of Mg^+^.

Several epidemiological studies have reported an inverse relationship between dietary Ca^2+^ intake and blood pressure and/or hypertension (52,77,78,81,86,89–94). Our model predictions are aligned with the observations from these studies. Our model simulations showed that Ca^+^ deficiency increased the total peripheral resistance but did not change cardiac output (Fig. 4). Additionally, our model predicted that dietary Mg^2+^ deficiency has a stronger effect on mean arterial pressure than dietary Ca^2+^ deficiency (Fig. 4). In fact, two cross-sectional studies have reported dietary Mg^2+^ intake to have a stronger association with blood pressure compared to dietary Ca^2+^ intake (81,95). Since calcitriol deficiency significantly lowers plasma [Ca^2+^], it has same effect as dietary Ca^2+^ deficiency on mean arterial pressure, i.e., an inverse association (54–56,96,97), though the impact is slightly lower.

Primary hyperparathyroidism has been associated with osteoporosis (65–67) and hypertension (60–64). Our model predictions suggested that the elevated plasma PTH, Ca^2+^, and calcitriol are primarily responsible for these two disorders (Fig. 5). These three factors overactivate the RAAS and increase vascular resistance, which in turn increases the mean arterial pressure. The elevated PTH, calcitriol, and AT1R-bound Ang II also increase bone loss (Fig. 5).

In summary, we have developed a computational model representing the interplay between Ca^2+^ and Mg^2+^ homeostasis and blood pressure regulation in a male rat. The model was used to understand the underlying mechanisms involved in (i) regulating Ca^2+^ and Mg^2+^ balance during different hypertensive stimuli, (ii) blood pressure regulation during dietary Mg^2+^ deficiency, dietary Ca^2+^ deficiency, and vitamin D_3_ deficiency, and (iii) Ca^2+^, Mg^2+^, and blood pressure regulation during primary hyperparathyroidism.

## Limitations of the study

The present model is based primarily on a male rat. However, there are many known sex differences in blood pressure regulation (29,98,99). Sex differences also occur in renal handling of Na^+^, Ca^2+^, and Mg^2+^ (100–103). A worthwhile extension would be to develop sex-specific models for Ca^2+^, Mg^2+^, and blood pressure regulation under various physiological and pathophysiological conditions.

The actions of the transporters and channels along the nephron cell membranes that regulate Na^+^, Ca^2+^, Mg^2+^, and fluid balance are represented implicitly in our model. A possible extension of the model would be to explicitly model these transporters and channels as done in epithelial transport models (101,102,104). The benefit of coupling individual nephron with whole kidney dynamics would be in simulating the administration of drugs that target these transporters. The exact action of the drug could be simulated instead of inferred.

The present model does not include potassium (K^+^) which is a key mineral in blood pressure regulation. Higher K^+^ levels have been associated with lower blood pressure (105–107). Its blood pressure-lowering effects stem from its capacity to induce vasodilation, a process mediated by vascular cell hyperpolarization (108). Additionally, K^+^ influences blood pressure by increasing Na^+^ excretion, modulating baroreceptor sensitivity, reducing vasoconstrictive sensitivity to norepinephrine and angiotensin II, and increasing Na-K-ATPase activity (108). A future extension of the model would be inclusion of K^+^ regulation (109,110) and its interaction with various components of blood pressure regulation.

## Data and code availability

The code generated in this study can be accessed at https://github.com/Pritha17/Blood-pressure-and-calcium-magnesium-homeostasis-regulation.

## Acknowledgement

This work was supported by the Canada 150 Research Chair program, National Sciences and Engineering Research Council of Canada (NSERC) Discovery Grant (RGPIN-2019-03916), and Canada Institutes of Health Research (CIHR) Project Grant (TNC-174963) to A.T.L.

## Author contributions

Conceptualization, P.D. and A.T.L.; Methodology, P.D. and A.T.L.; Software, Validation, Formal Analysis, and Investigation, P.D.; Resources, A.T.L.; Data Curation, P.D.; Writing – Original Draft, P.D. and A.T.L; Writing – Review & Editing, P.D. and A.T.L.; Visualization, P.D.; Supervision, P.D. and A.T.L.; Funding Acquisition, A.T.L.

